# A quantitative review of relationships between ecosystem services

**DOI:** 10.1101/017467

**Authors:** Heera Lee, Sven Lautenbach

**Author notes:** Corresponding author; *Email address:* (Heera Lee).

## Abstract

Ecosystems provide multiple ecosystem services to society. Ignoring the multi-functionality of land systems in natural resource management generates potentially trade-offs with respect to the provisioning of ecosystem services. Understanding relationships between ecosystem services can therefore help to minimize undesired trade-offs and enhance synergies. The research on relationships between ecosystem services has recently gained increasing attention in the scientific community. However, a synthesis on existing knowledge and knowledge gaps is missing so far. We analyzed 67 case studies that studied 476 pairwise combinations of ecosystem services. The relationships between these pairs of ecosystem services were classified into three categories: “trade-off”, “synergy” or “no-effect”. Most pairs of ecosystem services (74%) had a clear association with one category: the majority of case studies reported similar relationships for pairs of ecosystem services. A synergistic relationship was dominant between different regulating services and between different cultural services, whereas the relationship between regulating and provisioning services was trade-off dominated. Increases in cultural services did not influence provisioning services (”no-effect”). We further analyzed the pattern of relationships between ecosystem services across scales, land system archetypes and methods used to determine the relationship. Our analysis showed that the overall pattern of relationships between ecosystem services did not change significantly with scale and land system archetypes. However, some pairs of ecosystem services showed changes in relationships with scale. The choice of methods used to determine the relationship had an effect on the direction of the relationship: studies that employed correlation coefficients showed an increased probability to identify no-effect relationships, whereas descriptive methods had a higher probability of identifying trade-offs. The regional scale was the most commonly considered, and case studies were biased among different land system archetypes which might affect our ability to find the effect of scale or land system archetypes on the pattern of relationships. Our results provide helpful information of which services to include in ecosystem services assessments for the scientific community as well as for practitioners. Furthermore, they allow a first check if critical trade-offs have been considered in an analysis.

## 1. Introduction

Decision making on resource managements received worldwide attention in the past decades given the urgent need to preserve ecosystems and find a sustainable balance between long-term and short-term benefit and costs of human activities (Berkes and Folke, 1998; MA, 2005; Carpenter et al., 2009; Liu et al., 2015). However, a management decision can cause undesirable consequences if it lacks understanding of the complex nature of ecosystems which lead to the multi-functionality of land systems (Holling, 1996; Bennett et al., 2009). A land system does not provide only one function but combinations of a variety of overlapping functions (Bolliger et al., 2011, p.203), each of which provides different ecosystem goods and services to society. Land systems thus have a potential to provide multiple ecosystem services (ES) (Burkhard et al., 2009; Tallis and Polasky, 2009; Mastrangelo et al., 2014; Schindler et al., 2014). Due to functional trade-offs and synergies among the different components of this multi-functionality within the land, a decision potentially influences which services people can get or lose at the same time (Wiggering et al., 2006; Paracchini et al., 2011). Therefore, a comprehensive understanding of the multi-functional land system and of the different ES derived from it is crucial in natural resource management to avoid undesired and often unaware trade-offs and to enhance synergies among ES (Rodríguez et al., 2006; Hillebrand and Matthiessen, 2009; Bolliger et al., 2011; Mastrangelo et al., 2014). A key challenge that decision makers face now is to consider multiple ES and their potential consequences rather than focusing only on a few services in isolation (Cork et al., 2007; Tallis and Polasky, 2009).

The concept of multi-functionality has originally been developed at the landscape scale (Bolliger et al., 2011; Mastrangelo et al., 2014). However, it can be transferred to larger scales at which parts of the multi-functionality present at the landscape scale might be hidden due to aggregation effects. Likewise, the concept can be applied at smaller scales but one has to keep in mind that some functions might diminish at small scales such as functions that lead to water regulation, seed dispersal, pollination and pest control that connect different parts of the landscape. Therefore, interactions across multiple scales are important to be considered in decision-making.

The global research community endeavors to elaborate the concept of ES both in theory and practice to preserve multiple ES (MA, 2005; Carpenter et al., 2009). The Millennium Ecosystem Assessment (MA, 2005) has raised the awareness of the importance of identifying multiple ES and their interactions (Raudsepp-Hearne et al., 2010; Willemen et al., 2012). The number of publication has risen rapidly in last decades on this issue (Bennett et al., 2009). Bennett et al. (2009) stressed the importance of understanding direct and indirect relationships among multiple ES. Two recent review studies (Mouchet et al., 2014; Howe et al., 2014) addressed aspects of relationships between ES. Mouchet et al. (2014) provided a methodological guideline for assessing trade-offs between ES, whereas Howe et al. (2014) analyzed relationships between ES with a focus of beneficiaries and users. However, both studies did not analyze pairwise relationships between ES, which is a first step to investigate relationships among multiple ES (Chan et al., 2006; Raudsepp-Hearne et al., 2010; Jopke et al., 2014). Kandziora et al. (2013) provided a matrix of pairwise relationships between ES on a conceptual level, but the relationships between ES have not been studied so far based on case study results. In this study, we aim at filling this gap with a quantitative review of relationships between ES based on the published literature.

Recent studies focusing on multiple ES have taken several perspectives. The concept of “bundles” of ES has been commonly applied in the assessment of provisioning multiple ES in a landscape (e.g. Raudsepp-Hearne et al., 2010; Martín-López et al., 2013). This approach tries to identify groups of ES that co-occur repeatedly in landscapes showing patterns of the provision of ES derived from the different land use and land cover types (Raudsepp-Hearne et al., 2010; Turner et al., 2014). It is frequently based on a GIS analysis at the landscape or the regional scale (O’Farrell et al., 2010; Nemec and Raudsepp-Hearne, 2012). Often complementary statistical or descriptive analysis have been used to identify the bundles. Another research line tends to focus on ecosystem processes and functions that underpin ES (Dickie et al., 2011; Lavorel et al., 2011). The relationships among multiple ES are either identified by statistical analysis of field data or by the analysis of the output process models such as LPJ-GUESS (Smith et al., 2001) or SWAT (Arnold et al., 1999) see e.g. Lautenbach et al. (2013).

Relationships of ES pairs can be categorized into ‘trade-off’, ‘synergy’, and ‘no-effect’. The term ‘trade-off’ in ES research has been used when one service responds negatively to a change of another service (MA, 2005). An attempt to maximize the provision of a single service will lead to suboptimal results if the increase of one service happens directly or indirectly at the cost of another service (Holling, 1996; Rodríguez et al., 2006; Haase et al., 2012). When both services change positively in the same direction, the relationship between two ES is defined as synergistic (Haase et al., 2012) - this is often called also a ‘win-win’ relationship (Howe et al., 2014). When there is no interaction or no influence between two ES this is defined as a ‘no-effect’ relationship.

The relationship between a pair of ES can differ across different scales and across different socio-ecological systems (Kremen, 2005; Hein et al., 2006; Bennett et al., 2009). An example for this is the “externality” of a decision on a certain service as pointed out by Rodríguez et al. (2006): a decision that seems to influence ES positively for a specific region might cause substantial trade-offs in areas nearby or faraway (e.g. teleconnection) (Liu et al., 2013). If the effects of this decision are viewed at a larger scale including all those negatively influenced areas, the relationship between ES might be characterized by a trade-off. Cimon-Morin et al. (2013) showed in their review study that the relationship between biodiversity and ES changes with scale and region. The relationship between carbon storage and habitat was, for example, described mainly as synergistic at the global scale, but at a finer scale regions of high biodiversity and high carbon storage might be disjunct leading to a trade-off relationship. Furthermore, the relationship can change in different land systems. In other word, a decision on increasing a service can affect the other services differently in different locations. For example, Westa et al. (2010) showed differences in a trade-off relationship between carbon sequestration and food provisioning among regions.

Given the importance of understanding relationships between ES we conducted a quantitative review on relationships between pairs of ES based on case studies. We addressed three key hypotheses to investigate the relationships between ES. First, ES pairs show a preferred interaction and relationship with each other; second, this relationship is influenced by the scale at which the relationship had been studied as well as by the land system; and third, this relationship is further affected by the method applied to characterize the relationship.

## 2. Material and methods

### 2.1 Literature search

We carried out a literature search in the ISI Web of Knowledge database based on combinations of keywords including “ecosystem service*” or “environmental service*” or “ecological service*” in the first part, and “trade-off*” or “tradeoff*” or “synerg*” in the second part of the topic field. We limited the time period from 1998 to 2013, but decided to include four relevant studies published in 2014 in addition. Our query resulted in 585 scientific papers.

We only included case studies written in English. Studies that did not analyze the relationships between ES were clearly out of scope and therefore not further considered. If a case study analyzed more than one ES pair, we considered all pairwise combinations. In total our analysis is based on 67 case studies - with 476 ES pairs.

### 2.2 Database and classification

The ES categories were defined according to the Common International Classification of Ecosystem Services (CICES) classification V4.3 (Haines-Young and Potschin, 2013). CICES is one of the widely applied ES classification systems (e.g. Millennium Ecosystem Assessment (MA) (MA, 2005), National Ecosystem Services Classification System (NESCS) and Final Ecosystem Goods and Services Classification System (FEGS-CS) by the United States Environmental Protection Agency (Landers and Nahlik, 2013)), which has been also practically applied as a basis for the national ecosystem assessment, for example, in Belgium (Turkelboom et al., 2013), in Germany (Naturkapital Deutschland TEEB DE, 2014) and in Finland (Mononen et al., 2015). One of its advantages is that it contains a nested hierarchical structure (Haines-Young and Potschin, 2013). The highest level of CICES, the ‘Section’, distinguishes between provisioning, regulating and maintenance, and cultural services. The next hierarchical levels are ‘Division’, ‘Group’, and ‘Class’ (Fig 1). The analysis of this study was mainly based on the ‘Group’ level of CICES (Fig 1, see Supplementary table ST1 for the detailed list). From now on ‘ES’ refers to the ‘Group’ level of ES in CICES unless mentioned.

**Figure 1:**
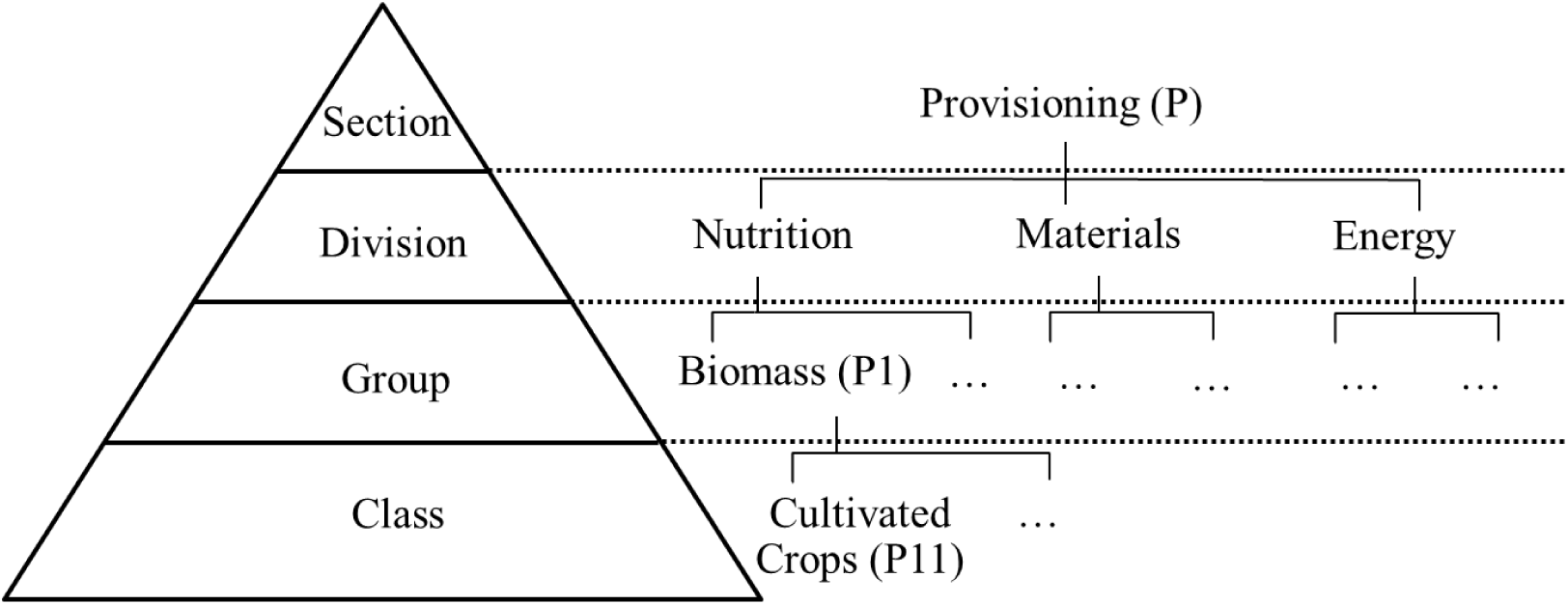
CICES nested hierarchy structure (left) and example of provisioning section and ES code in brackets (adapted from Haines-Young and Potschin (2013))

In this study, we focused on the pairwise relationships between ES as described in case studies. The relationship between each pair of ES was classified into three categories: “trade-off”, “synergy” and “no-effect”. “Trade-off” was assigned when one service increased with reduction of another service, whereas when both services interacted positively, “synergy” was assigned. When there was no interaction between two services, “no-effect” was assigned. If the direction of the relationship between the pair of ES was not clearly described, it was classified as “other”.

For case studies using correlation coefficients a threshold had to be defined to distinguish “no-effect” relationships from relevant relationships. There is no clear vote from the literature about such a threshold in the ES literature. Applied statistics textbooks agree that a Pearson’s correlation coefficient under 0.35 is characterizing either a negligible (Hinkel et al., 2003) or a weak relationship (Weber and Lamb, 1970; Mason and Lind, 1983; Taylor, 1990). In ES literature, however, a Pearson’s correlation coefficient of 0.2 is often considered as a meaningful correlation (e.g. Chan et al., 2006; Jopke et al., 2014). In this study, we assigned the “no-effect” label to relationships with a correlation coefficient between -0.25 and 0.25.

It was even more difficult for case studies using multivariate statistics to set a threshold to distinguish a “no-effect” relationship from a synergistic or trade-off relationship. The square of the factor loadings is the proportion of variance in each of the items (the observed traits) explained by the factor (the unobserved trait). For example, the factor loading of 0.32 is equivalent to 10% explained variance (Tabachnick and Fidell, 1989). In this study, we used a threshold: if the loading was reported and it was greater than 0.32, the relationship was identified according to the direction (+ for “synergy” or - for “trade-off”) over the different factors or PCs. When the loading was too small, “no-effect” was assigned. When the loading was not reported at all and only the bi-plot was reported from the study, the direction of variables (+ for “synergy” or - for “trade-off”) was considered for the relationship.

The dominant relationship for each pair of ES was determined based on the relative importance of each relationship category. The ratio of studies in the dominant relationship category (Eq. 1) was calculated across all case studies – the category with the highest ratio was assigned as the dominant relationship for each pair of ES. We used the term “level of agreement” to describe the certainty of relationships from the case studies.

The level of agreement of a pair of *ES*_*i*_ and *ES*_*j*_ is calculated as

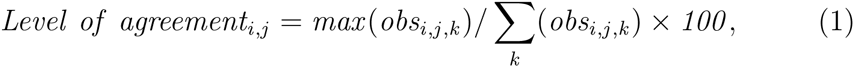

where *obs*_*i,j,k*_ is the number of observations for the pair of ES*i* and *j* in the relationship category *k*. The higher the level of agreement for a pair of ES, the higher the percentage of studies that showed the same direction of relationship. If there was a tie between two or three categories for a pair or if the level of agreement did not exceed 50%, we assigned the pair to the “not decided” category.

The spatial scale of the case study was determined following the criteria provided by Martínez-Harms and Balvanera (2012) (Table 1) according to the size of the study area. The land system in which a case study took place was assigned according to the map of land system archetype (LSA) of Václavík et al. (2013) that matched the location of the study site. The LSA is a classification schemes of land systems based socio-economic, ecological and land use intensity factors (Václavík et al., 2013). When several LSAs overlapped within a study area, the dominant LSA was assigned if it covered more than 50% of the study area. Otherwise, all LSAs were considered - at maximum three LSAs were assigned to one pair of ES.

**Table 1:** Criteria used for classification

| Criteria | Categories | Rationale | Reference |
| --- | --- | --- | --- |
| Spatial Scale | Patch | $10\text{-}10^2 \text{ km}^2$ | Martínez-Harms and Balvanera (2012) |
| | Local | $10^2\text{-}10^3 \text{ km}^2$ | |
| | Regional | $10^3\text{-}10^5 \text{ km}^2$ | |
| | National | $10^5\text{-}10^6 \text{ km}^2$ | |
| | Global<sup>a</sup> | $> 10^6 \text{ km}^2$ | |
| Archetype | LSA 1 | Forest systems in the tropics | Václavík et al. (2013) |
|  | LSA 2 | Degraded forest/crop land systems in the tropics |  |
|  | LSA 3 | Boreal systems of the western world |  |
|  | LSA 4 | Boreal systems of the eastern world |  |
|  | LSA 5 | High-density urban agglomerations |  |
|  | LSA 6 | Irrigated cropping systems with rice yield gap |  |
|  | LSA 7 | Extensive cropping systems |  |
|  | LSA 8 | Pastoral systems |  |
|  | LSA 9 | Irrigated cropping systems |  |
|  | LSA 10 | Intensive cropping systems |  |
|  | LSA 11 | Marginal lands in the developing world |  |
|  | LSA 12 | Barren lands in the developing world |  |
| Method | Descriptive | Qualitative description without any explicit quantitative measures |  |
|  | Correlation | Measures of the degree of statistical dependency between two variables such as Pearson’s correlation coefficient or Spearman’s rank correlation coefficient |  |
|  | Regression analysis | Regression analysis such as (generalized) linear models |  |
|  | Multivariate statistics | Analysis of pattern in multidimensional data without assuming a dependent variable such as PCA, cluster and factor analysis |  |
|  | Other | The relationship between ES was already built in the quantifying ES process |  |

<sup>a</sup> When a study considered a certain continent (e.g. Europe), we considered it as a continental scale.

We differentiated between the method used to quantify ES (preparation of the results) and the method used to identify the relationship between the ES (analysis of the results). We only considered the latter in the analysis. If, for example, a study used GIS modeling to quantify ES and described the relationship between ES - based on the GIS analysis - qualitatively, we categorized the method for this pair as “descriptive“. The method used to identify the relationship was categorized into five groups: “descriptive”, “correlation”, “regression analysis”, “multivariate statistics“, and “other” (Table 1).

### 2.3 Statistical Analysis

To test our hypotheses that the scale, the LSA as well as the method used affect the dominant relationship of ES, we applied two statistical analyses. In the first step we focused on the overall pattern of relationships between pairs of ES. In the second step we tested each pair of ES separately for effects of scale, the LSA and the method used. Subsets of the data were prepared for each category of scale, LSA and method (Table 1). The minimum number of case studies to participate in the comparison was set to 10 for each subset. We combined the national, the continental and the global scale into one category, “large scale“, due to the limited number of case studies in these categories. Among 12 LSAs, only three LSAs (i.e. “boreal systems of the western world” (LSA3), “extensive cropping systems” (LSA7), and “intensive cropping systems” (LSA10)) satisfied this threshold to participate in the comparison. The method used could not be performed for the overall pattern analysis due to the limited number of case studies in the categories.

In the first step we tested the null hypothesis that the overall structure of the relationships between ES pairs was independent of scale and LSA. To compare the outcomes of different subsets of scales and LSAs, a bootstrap approach (Efron and Tibshirani, 1994) was used. The subset membership was permuted at the case study level during the bootstrap because case studies often analyzed multiple pairs of ES leading non-independence between ES pairs from the same case study. For each bootstrap sample a measure of similarity between the original data and the permuted subset was calculated. As the measure of similarity, the Euclidean distance between the two subsets of ES relationships normalized by the total number of ES pairs in the subset was used. This allowed us to test the null hypothesis that both subsets belong to the same underlying distribution.

Afterwards, we tested each pair of ES separately for effects of scale, the LSA and the method based on the contingency table. After the contingency table for each pair was created, we fitted generalized linear model with a Poisson distribution for a model with the number of elements in a category as the response and the type of relationships and scales, LSAs or the methods as predictors. We tested for the significance of the differences of deviances by comparing the saturated model which contained the interaction between both factors to the model with just the main effects (Faraway, 2005). Since this analysis can only be applied for pairs studied at all scales or LSAs, we were only able to analyze 14 pairs of ES with respect to effects of scale. For the effect of LSA none of pairs was studied in all 12 LSAs. Therefore, we tested for the pairs which were studied in multiple LSAs: this led to the analysis of 19 ES pairs. The analysis of the effect of the method used to identify the type of relationship was done at the level of the case studies and not at the ES pair level since case studies typically applied the same method. We excluded the “other” category for the analysis. All analyses were performed using R version 3.2.0 (R Core Team, 2015).

## 3. Results and Discussions

### 3.1 Empirical pattern of the relationships between ecosystem services

Among the 48 types of ES defined at the class level in CICES, 33 - including one abiotic service (i.e. renewable abiotic energy source) - were found in our data set (Fig 1, Supplementary table ST1). The most studied ES class was “global climate regulation service” (n = 114) followed by “cultivated crops” (n = 103), “physical use of landscape” such as hiking (n = 93), and “maintaining nursery population and habitats” (n = 85). We found 207 different combinations of ES at the CICES class level (Fig 1). More than half of those combinations at the class level (n = 105) were, however, recorded only one time. Since this did not provide enough support to analyze patterns, we decided to drop the analysis at the class level. At the group level in CICES 94 types of combinations of ES pairs were analyzed (Fig 1, Supplementary table ST1). A pair of two ES that belonged to the same CICES group but to different CICES classes was analyzed as well. Figure 2 shows the empirical pattern of pairwise relationships between ES groups – non-empty cells at the main diagonal refer to pairs of ES classes that belong to the same CICES group. To the best of our knowledge, this is the first study in which such a comprehensive matrix of relationships between ES has been compiled based on case study results.

**Figure 2:**
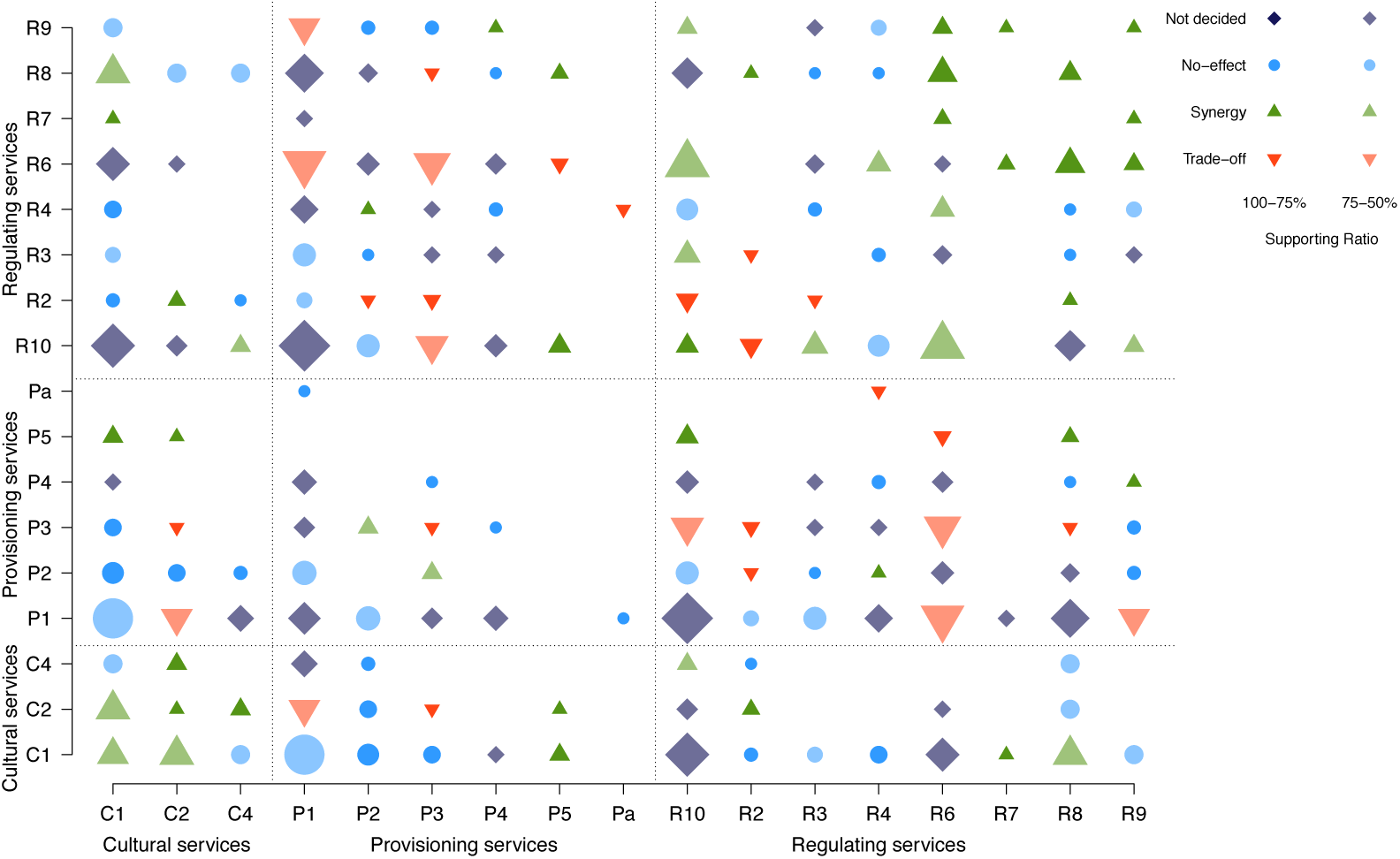
Result from analysis of 67 case studies with 476 pairs of ecosystem services, showing the empirical pattern of relationships between them. X and Y axis represent the ecosystem service classification code used in the analysis. The size of the symbol indicates the square root scaled number of studies. The color intensity represents the level of agreement. C: Cultural services, P: Provisioning services, R: Regulating services. C1: Physical and experiential interactions, C2: Intellectual and representative interactions, C4: Existence and bequest, P1: Biomass provisioning, P2: Water provisioning, P3: Materials for production and agricultural uses, P4: Water provisioning (i.e. non-drinking purpose), P5: Energy, Pa: Abiotic provisioning, R10: Atmospheric composition and climate regulation, R2: Mediation by ecosystems, R3: Mass flows regulation, R4: Liquid flows regulation, R6: Life cycle maintenance, habitat and gene pool protection, R7: Pest and disease control, R8: Soil formation and composition regulation, R9: Water condition

The number of observations available to identify the dominant relationship ranged between 1 and 29. Twenty-one types of pairs of ES at the group level were observed only one time and more than half of the pairs (n = 61) were supported by less than 5 observations. Only 12% of the pairs were supported by more than 10 observations. The most studied pair of ES at the group level was the pair “atmospheric composition and climate regulating” (R10) and “biomass provisioning” (P1) services with 29 observations.

The level of agreement ranged from 25% to 100% (Fig 3). For 74% of the pairs, the level of agreement to determine the dominant relationship was higher than 50% – the other pairs were assigned to the “not decided” category (n=24).

**Figure 3:**
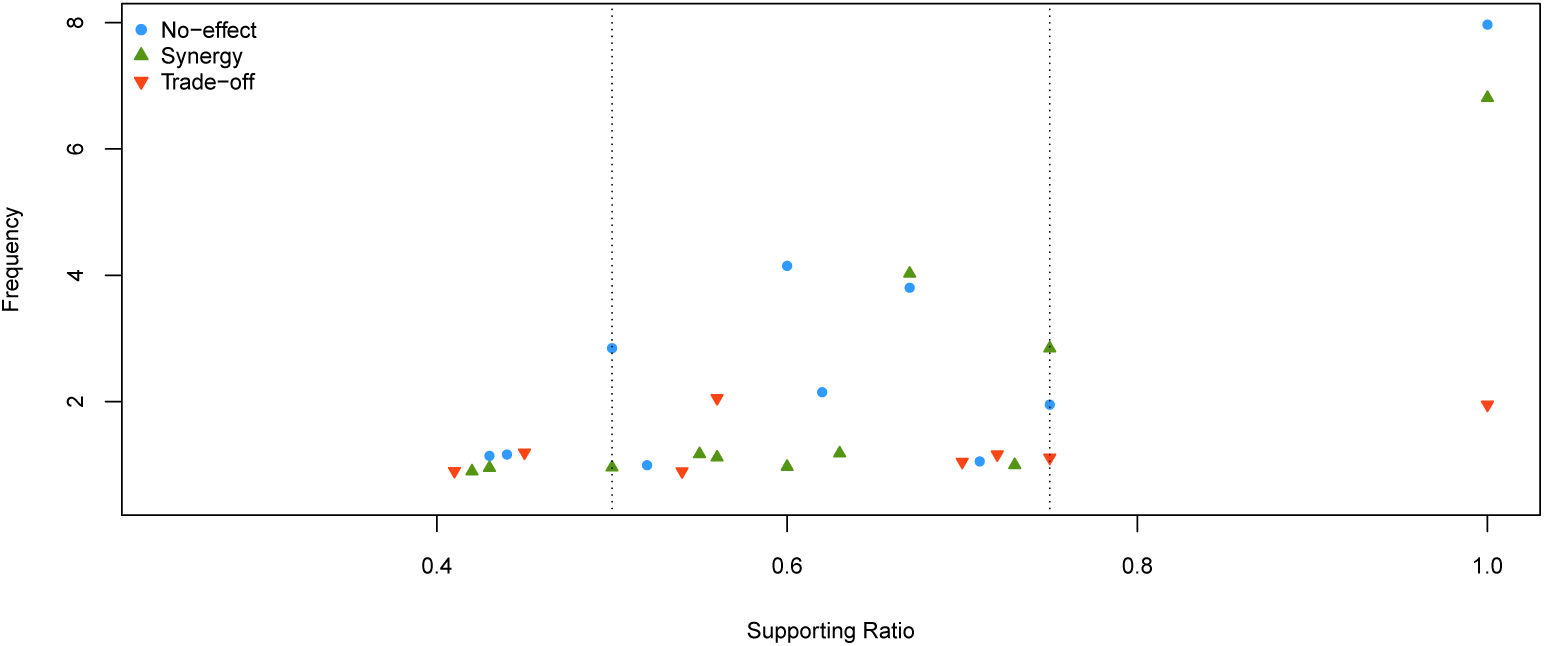
The distribution of the level of agreement (jittered for clarity) to determine the dominant relationship excluding pairs with a single observation. The shape of symbols indicates the dominant relationship.

The relationship between regulating services was dominated by a synergistic relationship, which means that regulating services are likely to increase if a management action increases other regulating services. On the other hand, provisioning services and regulating services tended to trade-offs (Fig 2), which means that when a provisioning service increases, a regulating service is likely to decrease and vice versa. Cultural services showed a trend for synergistic effects mainly with other cultural and regulating services, and a no-effect relationship with provisioning services. Note that this pattern of relationships shown here does not necessarily imply causality.

#### 3.1.1 Trade-off dominated relationships

The level of agreement for the trade-off relationships ranged between 54% and 100%. The most agreed trade-off relationship among those pairs with more than 5 observations was “biomass for production such as timber and fodder” (P3) and “atmospheric composition and climate regulation” (R10) with a level of agreement of 75% (n=8). On the one hand forests are important in terms of carbon fixation and storage, but on the other hand they are in many land systems used for timber production. In this case, a decision on how long forests are kept as carbon sinks or when trees are cut to be used for timber production generates trade-offs. Different forest management schemes influence the type of services from which people obtain benefits, which generates such trade-off among them (Backéus et al., 2005; Seidl et al., 2007; Olschewski et al., 2010).

A clear agreement on a trade-off relationship was also found for the relationship between the pair of “life cycle maintenance, habitat and gene pool protection” (R6) and “food provisioning” services (P1) with a level of agreement 72% (n=18). Previous studies pointed out a negative relationship between agricultural intensity and natural habitat (Mattison and Norris, 2005; Reidsma et al., 2006; Phalan et al., 2011). In order to compensate the loss of habitat in agricultural areas, more sustainable farming managements were often suggested (Altieri, 1999; Landis et al., 2000; Altieri, 2004; Lichtfouse et al., 2009) such as organic farming, which promises to increase ES nursery and habitat protection. However, there are doubts whether this allows producing sufficient food to feed the world population (Bengtsson et al., 2005; Zhang et al., 2007; de Ponti et al., 2012). Organic farming was found to increase species richness by providing better habitats and nursing ES (Bengtsson et al., 2005), but at the same time, meta-analyses showed that crop yield could be lowered by up to 20-34% compared to conventional farming (de Ponti et al., 2012; Seufert et al., 2012).

However, “pollination and seed dispersal” (R61) in the CICES class level, which belongs to “life cycle maintenance, habitat and gene pool protection” (R6) in the CICES group level, showed a synergistic relationship (e.g. Boreux et al., 2013). Overall 35% of the global production comes from crops that depend on animal pollinators (Klein et al., 2007), which might lead to a synergistic relationship between food provisioning and habitat protection (Aizen et al., 2008; Lautenbach et al., 2012; Garibaldi et al., 2013). It was not seen at the aggregated group level of CICES due to the limited number of case studies on R61.

#### 3.1.2 Synergy dominated relationships

The level of agreement for synergistic relationships varied between 55% and 100% (Fig. 3). The strongest synergistic relationship was found in the group of regulating services. Especially “habitat and gene pool protection services” (R6) showed a clear synergistic relationship with most other regulating services. Regulating services have been described as generally associated with ecosystem processes and functions (Kremen, 2005; Bennett et al., 2009; de Groot et al., 2010) and mostly positively related to biodiversity (Balvanera et al., 2006; Mace et al., 2012; Harrison et al., 2014). de Groot et al. (2002) defined “habitat and gene pool protection services” (R6) as a basis for other functions, which is in line with its observed synergistic relationship with other regulating services. The synergistic relationship between “habitat and gene pool protection services” (R6) and “soil formation regulating services” (R8) with a high level of agreement (88%) has been reported by studies that emphasized the interactions between soil functions and the role of soils in living habitats (e.g. Young and Ritz, 2000; Crawford et al., 2005; de Groot et al., 2010; Larvelle, 2012).

Another relatively strong synergistic relationship was found among the group of cultural services. Among pairs of cultural services, four out of five showed a dominant synergistic relationship. This is in line with findings from Daniel et al. (2012) on interrelationships between cultural service categories such as aesthetic services that contribute to the provisioning of recreation services, which leads to the synergistic relationship between them.

#### 3.1.3 No-effect dominated relationships

The level of agreement for no-effect relationships varied between 52% and 100%. The dominant no-effect relationship was found between provisioning and cultural services. Among pairs of provisioning and cultural services, “water provisioning service” (P2) and “physical and experimental interactions” (C1) was the most agreed no-effect relationship with a level of agreement of 100% (n = 7)

The dominant no-effect relationship between provisioning and cultural services could be explained by common drivers (Bennett et al., 2009) and different land use designs when the services occur in different locations (Raudsepp-Hearne et al., 2010). Bennett et al. (2009) proposed “common drivers” to understand relationships between ES. For example, introducing agricultural tourism by allowing people to watch the production process increases cultural services, but does not affect the amount of the agricultural production (Bennett et al., 2009, p.4). In this case, cultural and provisioning services do not share a common driver, therefore the relationship between them is no-effect. Another explanation for the no-effect relationship between provisioning and cultural services would be that cultural services such as tourism and cultural heritage are often captured in protected areas (e.g. national parks) where no production activity would be allowed (e.g. Martín-López et al., 2007; Raudsepp-Hearne et al., 2010). However, there was a disagreement on this relationship. Rodríguez et al. (2006) described the relationship between provisioning and cultural services as a ‘trade-off’ relationship – forest management for timber production could for example discourage recreational visits to this forest. It might depend on the types of ES whether they share a common driver or location to derive synergies or trade-offs.

Here we note that the types of cultural services that were covered in the analysis were rather limited; 69% of those case studies that analyzed cultural services focused on “physical and experimental interactions” (C1), whereas “spiritual services” (C3) were not considered at all in the studies analyzed.

#### 3.1.4 Sensitivity of the pattern towards changes in the threshold of the level of agreement

To determine the dominant relationship, we used 50% as a threshold for the level of agreement (Eq. 1) following majority rule. If the threshold was raised up to 70%, about 20% of pairs of ES were influenced by the decision and changed to the “not decided” category (see Table 2 and Fig 3). However, the overall direction of the dominant relationships between groups of ES (i.e. the “section” level of ES (Fig 1)) did not change thereby. See Supplementary Figure SF4 where we present the relationship matrix of pairs of ES with the threshold 70%.

**Table 2:** The number of pairs of ecosystem services in each category of relationships under the 50% and 70% threshold conditions.

|  | Trade-off | Synergy | No-effect | Not decided |
| --- | --- | --- | --- | --- |
| 50% threshold | 13 | 33 | 24 | 20 |
| 70% threshold | 10 | 24 | 16 | 40 |

#### 3.1.5 Sensitivity of the pattern towards the analysis at the CICES group level

Results might be potentially influenced by using the CICES group level for the analysis. However, we assume that only a single “not-decoded” pair has to be considered as an artifact from the aggregation of ES at the CICES group level (Fig 1): the pair of “physical and experiential interactions” (C1) and “soil formation and composition” (R8). While most case studies for this pair were conducted at the same scale and in the same LSA using the same methodology, the direction of the relationship was different across the case studies. Six observations were synergistic, whereas five observations were identified as no-effect. All no-effect relationships were observed in “physical activities such as hiking and leisure fishing” (C12), whereas four among six synergy relationships were observed in “experiential use such as bird watching” (C11) at the class level in CICES (Fig 1).

Except this one case it was not possible to use the class level of CICES for the analysis due to the limited number of observations at this level. Our analysis at the group level in CICES provides an overall pattern of relationships over 94 pairs of ES. Furthermore, to our knowledge, the analysis of relationships between ES at the group level was rarely done before. Previous review studies provided results at a section level in CICES (e.g. provisioning, regulating, cultural services) (Rodríguez et al., 2006), or based only on examples (Bennett et al., 2009).

### 3.2 Scale and land system archetypes of ecosystem service pairs

The bootstrap approach did not reveal any significant difference between subgroups of the case studies based on scale or LSA. Neither spatial scale nor LSA membership had a significant influence on the overall pattern of the relationships between the services – p-values for each test are given in the Supplementary table ST3 and ST4.

The spatial scale of the studies was spread unevenly. The regional scale was most frequently studied (38%), followed by the plot scale (22%) and the continental scale (10%). The global scale was the least studied (6%) (Supplementary figure SF2). Forty-one pairs of ES (44%) were studied only at a single type of scale, which hindered the comparison of the relationship pattern among scales.

Among the 14 pairs of ES that were included in the contingency analysis, significant differences across scale were only identified for two pairs of ES: the pair of “soil formation and composition regulation” (R8) and “biomass provisioning” (P1) (p = 0.0067) and the pair of “soil formation and composition regulation” (R8) and “atmospheric composition and climate regulation” (R10) (p = 0.0321). Both pairs included “soil formation and composition regulation” (R8). The result for both pairs showed a synergistic relationship at the small scale, whereas at the larger scale, the relationship was no-effect and not-decided for the pair of R8 and P1, and the pair of R8 and R10, respectively. “Soil formation and composition regulation ” (R8) are generally considered not only as a direct driver for enhancing “biomass provisioning” (P1) in agricultural lands (Hobbs et al., 2008), but also as an indirect driver for enhancing carbon and nutrient cycling which can influence “atmospheric composition and climate regulation” (R10) by affecting biotic processes (van Breemen, 1993; Barrios, 2007). This synergistic role of “soil formation and composition regulation” (R8) was often studied in experiments at a finer scale (Six et al., 2000; Hobbs et al., 2008). At a larger scale, this relationship did not clearly appear – e.g. Jopke et al. (2014) showed a no-effect relationship at the continental scale.

In addition, there was only one pair which was considered at every scale. The results at each scale showed different relationships but the result was not statistically significant (p = 0.4213). It was the pair of “atmospheric composition and climate regulation” (R10) and “biomass provisioning” (P1): at the small scale (i.e. the plot, local scale) the dominant result was synergy (50%; n=3), whereas it was trade-off (54%; n=6) at the regional scale and no-effect (46%; n=5) at the large scale (national, continental and global).

Case studies were also unevenly distributed across LSAs (Supplementary figure SF3): only three types of LSAs (i.e. “boreal systems of the western world” (LSA3), “extensive cropping systems” (LSA7), and “intensive cropping systems” (LSA10)) among 12 were studied in more than 10 case studies. A geographical bias of the distribution of ES case studies was already stressed by Seppelt et al. (2011). The land system “boreal systems of the eastern world” (LSA4), “high-density urban agglomerations” (LSA5), and “irrigated cropping systems with rice yield gap” (LSA6) were not at all considered in the case studies. Thirty-two pairs of ES (34%) were studied at a single LSA. LSA10 was most frequently observed when only a single type of LSA was considered. At maximum, seven LSAs were considered for a pair of ES, the pair of “atmospheric composition and climate regulation” (R10) and “life cycle maintenance, habitat and gene pool protection” (R6).

While the overall pattern of relationships between ES at the group level was indifferent to LSA, a few ES pairs showed interesting differences across LSAs from the contingency analysis. The relationship for the pair of “life cycle maintenance, habitat and gene pool protection” (R6) and “atmospheric composition and climate regulating” (R10) showed significantly different across the LSAs (p = 0.0269): synergy in “forest systems in the tropics” (LSA1), “extensive cropping systems” (LSA7), and “intensive cropping systems” (LSA10), no-effect in “irrigated cropping system” (LSA9) and “marginal lands in the developed world” (LSA11), and not decided in “boreal systems of the western world” (LSA3). Stored carbon in vegetation and soil was generally measured to quantify climate regulating services (R10) in every LSA. However, for “habitat protection services” (R6) different approaches were used in different LSAs. A possible explanation is that in “forest systems in the tropics” (LSA1) and “extensive cropping system” (LSA7) species richness as well as carbon sequestration are positively influenced by the presence of forest instead of arable land areas, while in “irrigated cropping system” (LSA9) and “marginal land” (LSA11) such a clear common driver is missing. Other pairs which differed significantly across LSAs were the pair of “existence and bequest” (C4) and “biomass provisioning” (P1) (p = 0.0152) and the pair of “biomass provisioning” (P1) and “Water provisioning (i.e. non-drinking purpose) ” (P4) (p = 0.0331).

### 3.3. Methods used to determine the relationship

The results from the difference of deviance test showed that the influence of the choice of methods applied on the direction of the results was marginally significant (p = 0.0294). Correlation coefficient methods showed a higher probability to identify a no-effect relationship, whereas descriptive methods showed a higher probability to identify a trade-off relationship and less no-effect relationships. Multivariate statistics showed less no-effect relationships (Fig 5 and Fig 4).

**Figure 4:**
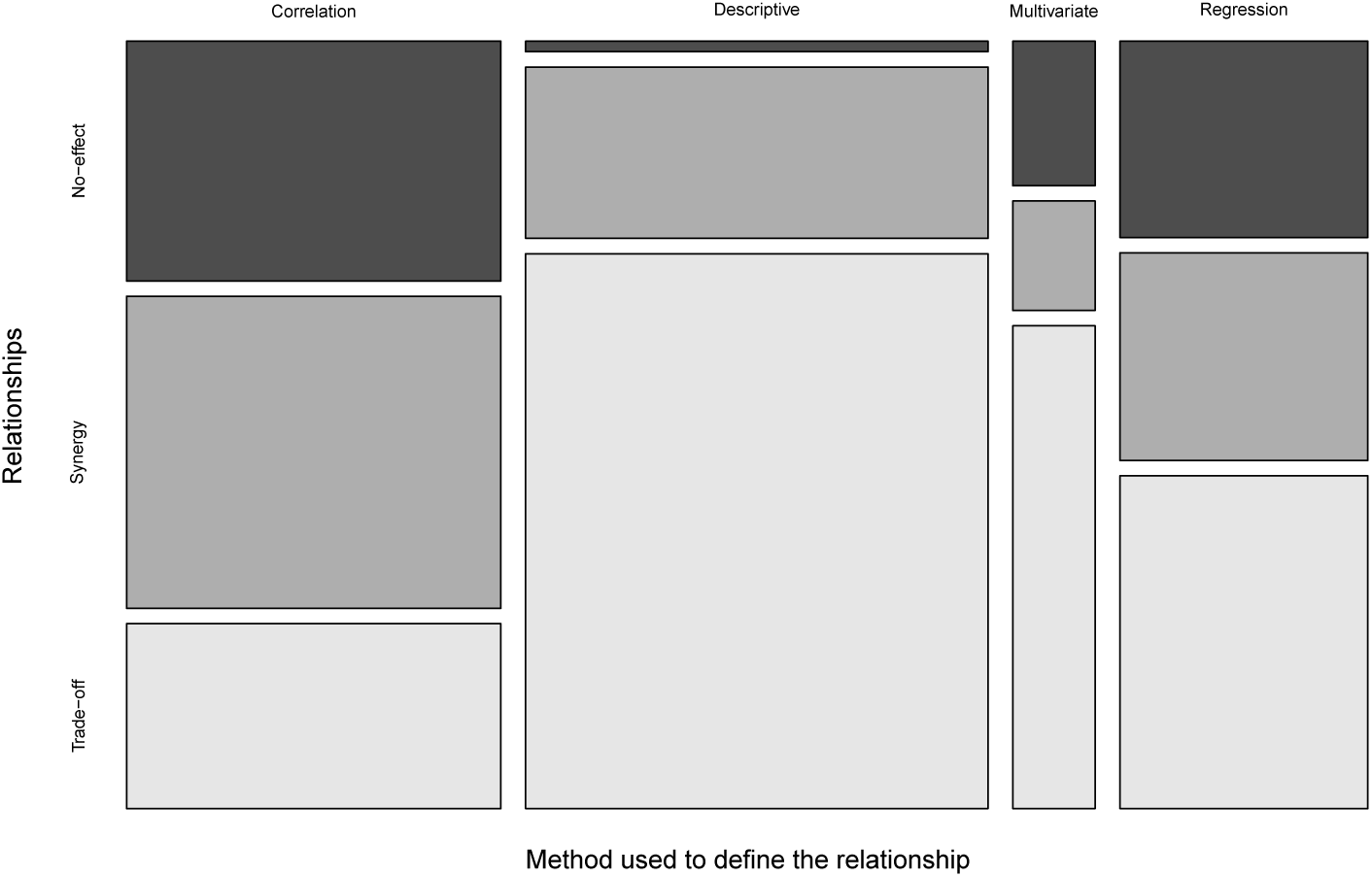
Mosaic plot for method used and the relationships between two ecosystem services

**Figure 5:**
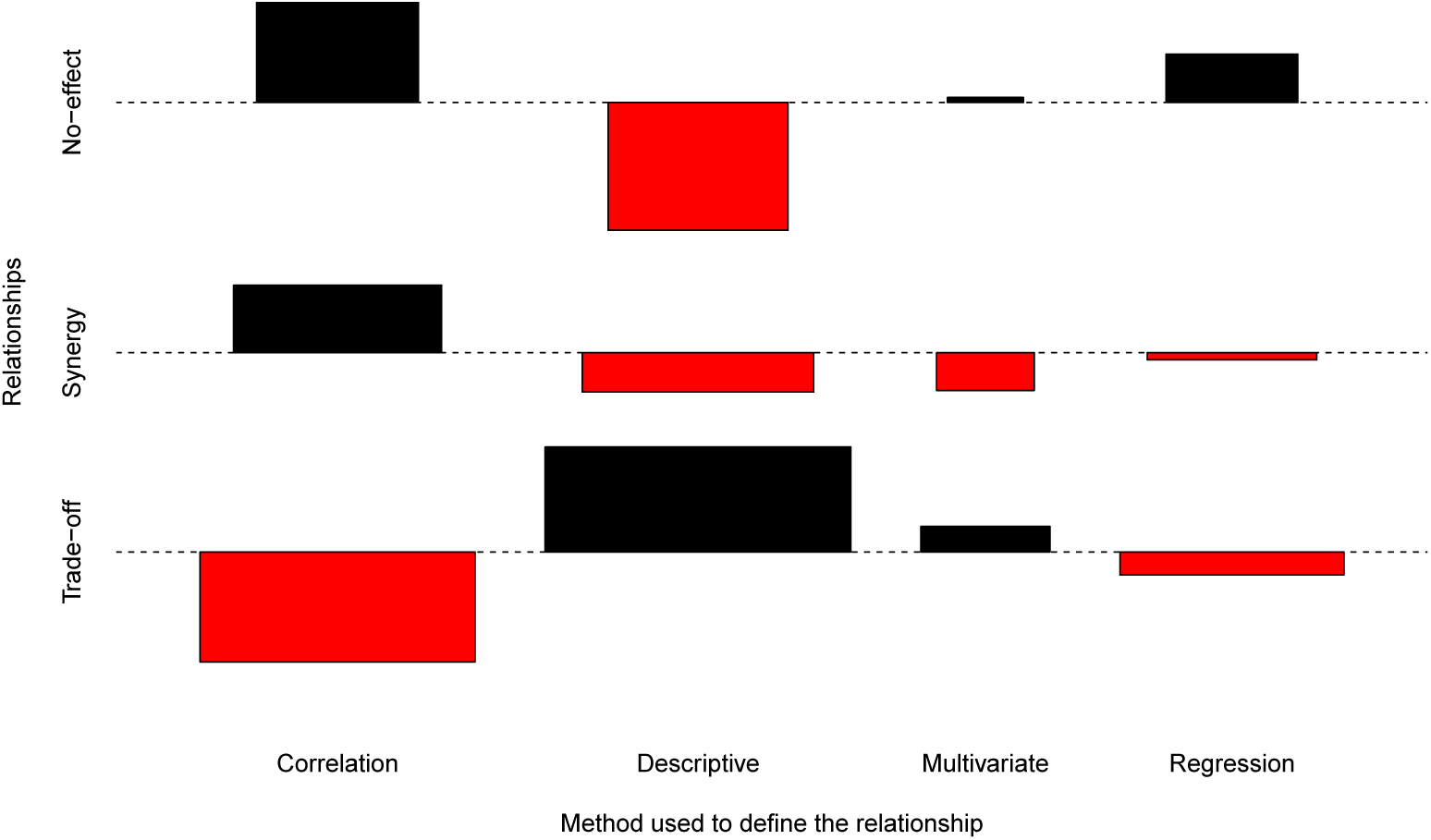
Association plot for method used in the relationships between two ecosystem services

It was problematic for case studies using multivariate statistics to set a threshold to distinguish “no-effect”. While it is possible to identify thresholds for the strength of the relationship based on the loadings in PCA or factor analysis as well as for the uncertainty of assigning an ES to a cluster, this was rarely done in practice. Multivariate statistics were frequently applied in trade-off of ES researches to identify bundles of ES by using PCA or factor analysis in order to find ES that tend to occur together (e.g. Lavorel et al., 2011; Maes et al., 2012). However, without an agreed threshold within ES research communities, using multivariate statistics to define relationships between ES might lead to ignorance of no-effect relationship. Since the assignment of ES to different bundles does typically neither include the strength of the association nor the attached uncertainty, no-effect relationships might be undetectable by the approach. Correlation approaches make it easier to define no-effect relationships based on the absolute strength of the correlation. If the correlation is stronger than a threshold, significance of the correlation should be tested – potentially corrected for nuisances such as spatial auto-correlation (Dormann et al., 2007).

Regression type I models were frequently used to describe the relationship between ES. From a theoretical point of view, the use of a regression type I model seems questionable to describe relationships between ecosystem services since the approach distinguishes ecosystem services into dependent and independent variables - errors are only considered for the dependent variable not for the independent variables. Only regression type II models (Legendre and Legendre, 2003) - which have not been used in the case studies-, in which errors for both predictors and response are considered seem appropriate to model ES relationships.

Methods were evenly distributed across the types of pairs of ecosystem services and across the scales. In other word, the decision on which types of method to use to define the relationship was neither influenced by the type of ecosystem services nor by the scale of the study.

It has been already reported (Vatn and Bromley, 1994; Jacobs, 1997; Martín-López et al., 2013) that the choice of the method to value ES can bias results. We emphasize here that not only valuation methods but also method used to define relationships should be chosen with a care. Researchers should be aware that their decision on methods used might limit the result in a certain direction.

### 3.4 Further limitations

Although our review was comprehensive and thoroughly conducted, we imposed constraints on our review that might have biased our result. We only considered peer-reviewed scientific articles written in English found in Web of Knowledge for our analysis. This might have excluded some pairs of ES that are only considered for a certain region in gray literature. However, using non peer-reviewed literature has the drawback that quality standards are lower (Pullin and Stewart, 2006; Nieto-Romero et al., 2014; Harrison et al., 2014).

## 4. Conclusions

Comprehensive information is required for well-informed policy decisions which do not ignore side-effects in multi-functional land systems. However, this information is often expensive and difficult to obtain. The missing information can directly and indirectly influence the policy decision as well as its impact on multi-functional land systems, and therefore human well-being (OECD, 2003). The fundamental challenge in practice is to minimize the inefficient and inappropriate impacts on provisioning of multiple ES by enhancing understanding of multi-relationships between ES. Making this information more explicit and accessible is more likely to drive at more balanced conditions (Carpenter et al., 2009).

In this study, we tried to fill the knowledge gap on relationships between ES by a synthesis of relationships between ES studied in published case studies. We identified typical relationships between a number of pairs of ES. To the best of our knowledge, this is the first study in which such a comprehensive matrix of relationships between ES has been complied. Our results provide an overview of relationships of ES studied so far together with the information on the level of agreement between study results. This equips practitioners with a practical summary to examine the underlying impacts of their decision in advance. Furthermore, our results might highlight pairs of ES for which more input is needed from the scientific community. The results might help further during the design of research programs and give important hints for decision makers and reviewers to check research plans and to ask critical questions with respect to research outcomes. If important relationships between ES could not be studied, our analysis might provide hints on the direction of the neglected effect.

While we were able to show that for a few pairs of ES the dominant relationship changed as a function of scale or of land system, we were not able to show this for the majority of cases. The limited number of case studies and the uneven distribution across ES groups, scales and land system archetypes is a potential explanation for it. Therefore, we encourage the development of a research agenda that allows filling those gaps to come to a more complete picture on relationships between different ES. Being able to predict the direction of a relationship between ES as a function of scale and land system would be an important step for decision support and ecosystem management but it would be by no means the end of the research agenda. We need higher quality studies that follow good modeling practice or analyze their data properly, reporting uncertainties along with point estimates, more evenly spread across the scales and land systems which reports not only the direction but also the strength of the relationship in a comparable way. Bundle analysis based on an overlay of relatively simple GIS tools presumably would not fulfill high quality standards and should be therefore treated with care. Based on the results of such data, a next step would be the performance of a meta-analysis to untangle more details on ES relationships.

## Acknowledgements

This project was funded by the EU FP-7 project OPERAs (grant number 308393). We acknowledge stimulating discussions with our project partners during the project meetings. We would like to thank Kimberly A. Nicholas, Stefan Schmidt, Sina Berger and Daniela Braun for valuable comments on an earlier version of the manuscript.

